# Deep Representation Learning Reveals Factorized Neural Codes in Whole-Brain Population Dynamics

**DOI:** 10.64898/2026.05.12.724368

**Authors:** Amina Abdelbaki, Paul Bandow, Karen Y. Cheng, Ilona C. Grunwald Kadow, Martin Paul Nawrot, Vahid Rostami

## Abstract

Extracting interpretable representations from high-dimensional whole-brain neural dynamics remains a major computational challenge in systems neuroscience. Here, we develop a wiring-agnostic deep-learning framework that combines convolutional encoding with temporal transformers to learn compact representations directly from volumetric calcium imaging of the entire *Drosophila melanogaster* brain, without neuronal identification or anatomical annotation. Applied to brain-wide activity recorded across 16 factorially combined sensory and internal-state conditions, the learned representations revealed a factorized organization of whole-brain dynamics: metabolic state, sensory modality, and stimulus valence emerged along three near-orthogonal axes from a classification objective based only on flat class labels and without explicit disentanglement constraints. Spatial attribution and brain region-level ablation analyses linked modality representations to anatomically distinct circuits, whereas state- and valence-related information showed a broadly distributed organization. Our approach provides a scalable framework for discovering interpretable representations of whole-brain neural dynamics.

## Introduction

Brains continually transform a torrent of external and internal sensory signals conditioned on the internal physiological state into coherent, adaptive behaviour. Although the population activity of even small nervous systems spans an extremely high-dimensional space, mounting evidence shows that these dynamics are often confined to low-dimensional manifolds that succinctly capture the essence of neural and behavioural states [1–11]. Such population-level structure suggests that the brain may compress information into behaviourally relevant coordinates, enabling efficient computation and generalization.

Complementary theoretical frameworks argue that brain states may correspond to structured patterns of neuronal population activity arising in recurrent circuits, where network dynamics settle into stable or slowly drifting configurations despite ongoing noise and fluctuating inputs [12–15]. Attractor-based computation provides one influential framework for understanding how such structured population states can support information processing. A rich taxonomy of attractor motifs, from continuous and discrete attractors to metastable multi-attractor landscapes, has been extensively studied theoretically and identified in combined experimental-theoretical studies across species and brain regions [14–22], suggesting that attractor computation is a general organizational principle of neural circuits. Yet whether geometric signatures consistent with such structured, low-dimensional dynamics can be recovered from mesoscale recordings spanning an entire brain, without knowledge of circuit identity or connectivity, remains an open question.

Concurrently, technical advances in volumetric imaging methods such as light-field microscopy now enable nearly complete brain-wide activity monitoring [23–29], capturing tens of thousands of neurons at several volumes per second. The fruit fly *Drosophila melanogaster* is particularly well-suited to exploit these capabilities: its compact brain fits entirely within the imaging volume, its genetic toolkit allows calcium indicators to be expressed pan-neuronally or in specific neuronal populations, and its neural circuits have been extensively mapped at the connectomic level [30–34]. Together, these properties raise the question whether the structured, low-dimensional dynamics so far characterized within individual brain regions also organize population activity at the scale of an entire brain — and, if so, whether such global structure is sufficiently rich to encode the fly’s sensory and internal context.

The fly’s chemosensory system offers a particularly compelling entry point, as odor and taste processing engage distributed circuits that are dynamically tuned by metabolic need. This state-dependent gating relies heavily on localized neuromodulation across distinct sensory hubs. For instance, pan-neuronal functional imaging has shown that metabolic state reshapes sensorimotor processing across the subesophageal zone [35], the primary taste processing center [36]. Similarly, hunger recruits diverse neuromodulatory pathways within the antennal lobe to selectively alter primary olfactory representations [37–41]. Further downstream in the central insect brain, dopaminergic neurons innervating the mushroom body integrate stimulus valence with physiological state [42], thereby reconfiguring associative networks to bias behavioral choice [43–47]. Although these studies demonstrate that internal states modulate chemosensory processing at multiple levels, their focus on isolated circuits leaves open whether metabolic state reorganizes information through local circuit modulation or through a distributed, brain-wide population code [48].

While these questions are now experimentally tractable, our ability to collect high-dimensional, brain-wide data has outpaced our ability to interpret it. Established dimensionality reduction methods — both linear approaches such as PCA [1], task-informed demixing approaches such as demixed PCA [49], and nonlinear techniques including latent variable models and contrastive embedding frameworks [50, 51] — have proven valuable for uncovering low-dimensional structure in neural populations, yet generally operate on pre-extracted neural time series, whether spike trains or segmented calcium traces, and may not fully exploit the multi-scale spatiotemporal structure present in volumetric recordings. Deep learning architectures trained directly on the imaging volumes offer an alternative: convolutional networks extract hierarchical spatial features [52–54], whereas transformer networks employing self-attention capture long-range temporal dependencies [55]. These complementary strengths offer a path toward data-driven discovery of interpretable structure in representations of whole-brain population dynamics without requiring prior cell segmentation, anatomical annotation, or region-of-interest selection.

Here we present a wiring-agnostic deep learning framework that couples a convolutional encoder with a temporal transformer to distill volumetric light-field calcium recordings of the entire adult *Drosophila* brain into a compact, functionally interpretable latent space. Applied to a dataset that factorially combines metabolic state, sensory modality and valence across 16 conditions, the framework learns representations directly from imaging volumes without information about neuronal identities, anatomical regions, or synaptic connectivity. The learned latent space organizes whole-brain population activity into condition-specific clusters whose geometry is factorized, with metabolic state, sensory modality, and stimulus valence emerging along approximately independent axes despite the absence of explicit disentanglement constraints. A linear classification head trained jointly with the encoder decodes each combination of metabolic state and stimulus condition with high accuracy on held-out data, demonstrating that mesoscale neural dynamics alone carry sufficient information to distinguish the fly’s physiological and sensory context. More broadly, these results provide direct evidence that the low-dimensional organization and state-dependent separation of neural population activity previously identified within individual circuits extend to the scale of an entire *Drosophila* brain.

## Results

We recorded whole-brain calcium activity by light-field microscopy in 82 head-fixed *Drosophila melanogaster* during controlled chemosensory stimulation (164 recordings; 71,423 input sequences; Fig. 1a–c; *Materials and Methods*). The experimental design combined three factors: metabolic state (fed vs. starved animals), sensory modality of stimuli (olfaction, gustation, or combined), and stimulus valence (appetitive, aversive, or conflicting), yielding 16 distinct experimental conditions (Fig. 1c). Volumes following stimulus onset were downsampled from the native 10 Hz to 1 Hz by retaining every tenth frame, reducing frame-to-frame temporal redundancy while capturing the slower dynamics of the evoked response. The resulting frames were collapsed to 2D mean-z projections, and arranged into short temporal sequences that capture the temporal evolution of the evoked response (Fig. 1b).

**Figure 1:**
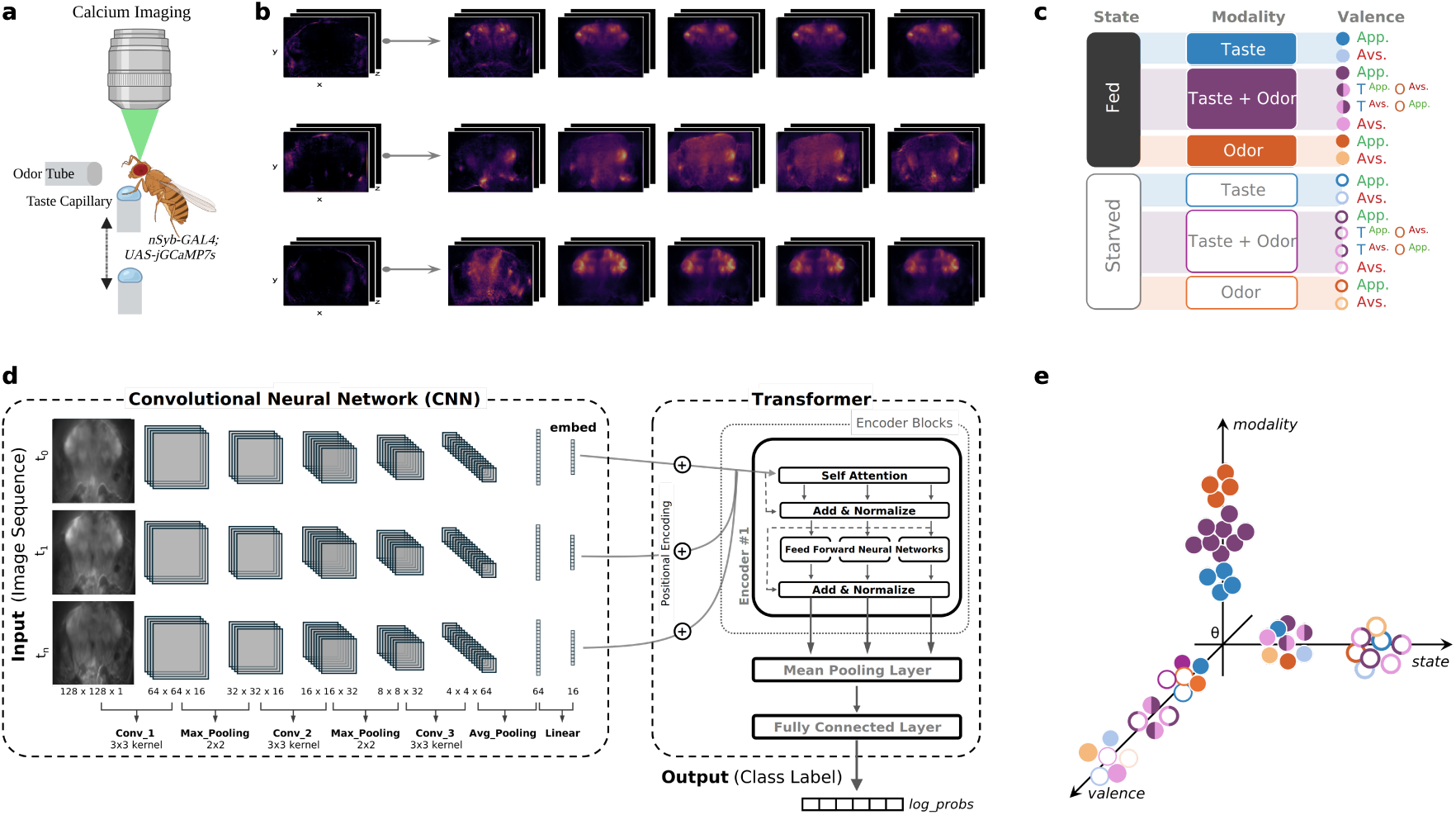
Experimental design and deep representation-learning framework. **(a)** Whole-brain calcium imaging in head-fixed *Drosophila melanogaster* expressing GCaMP pan-neuronally during controlled olfactory and gustatory stimulation. **(b)** Representative whole-brain image sequences used as model input; a pre-stimulus frame is shown for comparison. **(c)** Experimental design combining metabolic state (fed, starved), sensory modality (odor, taste, odor+taste), and stimulus valence (appetitive, aversive, or conflicting) across 16 conditions. **(d)** CNN–Transformer architecture. Individual image frames are spatially encoded by a convolutional neural network; frame embeddings are then processed by a temporal Transformer with self-attention and passed to a linear classification head. **(e)** Schematic illustration of the latent-space organization examined in this study, with dimensions corresponding to state, modality, and valence. Colours and symbols denote the experimental conditions shown in c.

To decode experimental conditions directly from these frame sequences, we trained a hybrid CNN–Transformer architecture (Fig. 1d). Each frame was passed through the convolutional encoder to obtain a compact embedding. The resulting sequence of frame-level embeddings was augmented with positional encodings and processed by a transformer encoder, which modeled dependencies across frames. A linear classification head then mapped the learned sequence representation to experimental condition. Successful classification therefore requires the upstream encoder to produce representations that are linearly separable at the classifier input. The objective constrains only this separability, not how the classes are arranged relative to one another; no explicit latent-space regularization was applied (Fig. 1e).

### The learned latent space organizes whole-brain activity along biologically meaningful axes

To assess what structure emerges in the learned representations, we organized the analysis around three classification tasks of increasing complexity (Fig. 1c): metabolic state alone (2-class), state and modality (6-class), and state, modality and valence (16-class). After training, sequence-level embeddings were extracted from the held-out test set and visualized using t-SNE (Fig. 2a–b). A shuffled-label control, trained with identical architecture and procedure but with class assignments randomly permuted at the recording level, served as a baseline to confirm that any observed structure reflects genuine stimulus and state-related information rather than artifacts tied to individual recordings. The accuracy achieved at the chosen embedding dimensionality (starred in Fig. 2d) substantially exceeded the shuffled control across all three tasks, establishing that the geometric structure visible in the latent space carries genuine decodable information (see *Materials and Methods*).

**Figure 2:**
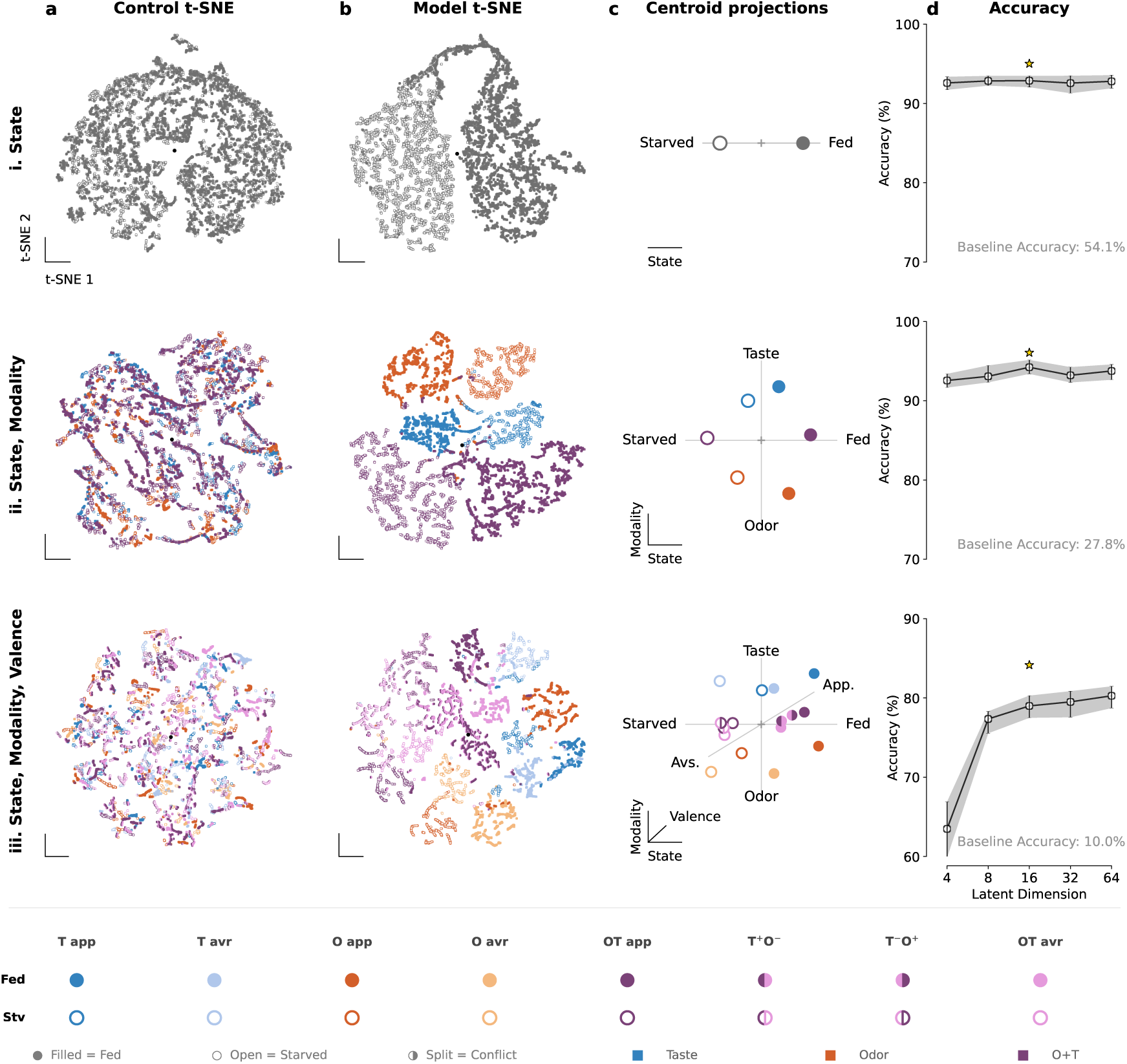
Learned latent representations reveal near-orthogonal axes for internal state, stimulus modality and valence. Rows show classification tasks of increasing complexity: (i) metabolic state (fed versus starved; 2 classes), (ii) state and sensory modality (odor, taste, odor+taste; 6 classes), and (iii) state, modality and valence (appetitive, aversive, or conflicting; 16 classes). **(a, b)** t-SNE projections of sequence-level embeddings from **(a)** shuffled-label controls and **(b)** trained models. **(c)** Class centroids in the 16-dimensional model latent space projected onto factor axes defined by group-mean contrasts for state, modality, and valence. No orthogonalization was applied. State and modality axes were near-orthogonal in the 6-class task (cos *α*_state, mod_ = *−*0.03); all three pairwise axis combinations were near-orthogonal in the 16-class task (cos *α*_state, mod_ = 0.06, cos *α*_state, val_ = 0.05, cos *α*_mod, val_ = *−*0.11). **(d)** Test accuracy as a function of latent dimensionality (*d*_model_ = 4, 8, 16, 32, 64). Stars indicate the dimensionality selected for the representations shown in a–c. Shuffled-label accuracy is shown for reference.

At the simplest level, embeddings from the 2-class metabolic state task formed two separated clusters in the t-SNE projection corresponding to fed (filled circles) and starved flies (open circles), while the shuffled control showed no comparable organization (Fig. 2i, a–b). As task complexity increased to state and modality, the latent space partitioned first by modality, with olfactory, gustatory, and combined conditions occupying well-separated regions, and then by metabolic state within each modality (Fig. 2ii, b).

In the full 16-class task, the t-SNE revealed a three-level hierarchical organization (Fig. 2iii, b). At the coarsest scale, embeddings separated by modality: odor classes clustered together, taste classes occupied a distinct region, and combined classes formed a third neighborhood, with one taste class positioned closer to the odor cluster. Within each modality cluster, a secondary separation distinguished metabolic states, with starved conditions systematically offset from fed conditions. At the finest scale, valence sub-classes were distinguishable within each state–modality sub-cluster, with appetitive and aversive conditions occupying distinct positions.

### Factor-axis projection reveals a near-orthogonal, factorized code

To move beyond the qualitative two-dimensional t-SNE visualization and quantify the geometric relationships among experimental factors in the native latent space, we defined a factor axis for each of the three experimental variables (state, modality, and valence) directly in the 16-dimensional embedding space (see *Materials and Methods*). Each axis was obtained as the difference between the group-mean embeddings of the corresponding factor levels (e.g., mean of all starved embeddings minus mean of all fed embeddings for the state axis; for factors with more than two levels, axes were defined from balanced combinations of level means). Class centroids were then projected onto these axes (Fig. 2c). Crucially, no orthogonalization was applied: the factor axes were used as computed from the data, and any observed orthogonality is therefore an emergent property of the learned representation rather than an imposed constraint.

In the 2-class task, the fed–starved contrast defined a single state axis, as expected for a binary centroid geometry, with all between-centroid variance lying along this one-dimensional subspace (Fig. 2ci). For the 6-class task, the state axis and modality axis were near-orthogonal (cos α_state,mod_ = −0.03), indicating that the network encodes these two factors along largely independent axes. Projection onto the two-dimensional plane spanned by these two axes captured 38.4% of total centroid variance, with the three modalities separated along one axis and fed-starved conditions offset along the other (Fig. 2cii). The remaining variance lies in directions that no single factor defines, such as differences between individual classes sharing the same state and modality. Projected onto each axis separately, every factor separates while the other collapse (Fig. S3a): fed and starved centroids fall on opposite sides of the state axis by comparable margins for all three modalities, and odor and taste centroids occupy opposite ends of the modality axis irrespective of state, with bimodal (*O* + *T*) conditions intermediate.

In the full 16-class task, all three pairwise angles between factor axes remained close to orthogonal (cos α_state,mod_ = 0.06; cos α_state,val_ = 0.05; cos α_mod,val_ = −0.11), indicating that state, modality, and valence are arranged along largely independent directions. Projection onto these three axes captured 24.4% of centroid variance and revealed the same hierarchical organization visible in t-SNE, now expressed in a coordinate system whose axes have explicit factorial meaning (Fig. 2ciii). Modality elicits the coarsest separation, metabolic state an intermediate offset, and valence the finest displacement. The same invariance holds when all three factors vary (Fig. S3b): each axis orders the centroids by its own factor with displacements of similar magnitude across the levels of the other two, appetitive and aversive conditions falling symmetrically on either side of the valence axis and the conflicting-valence classes (*T* ^+^*O^−^, T ^−^O*+) near its origin. The pairwise projections (Fig. S3c) show the corresponding two-dimensional arrangements, in which a centroid’s position along one axis is largely unchanged by its level on the other. Together, the latent geometry reveals that the CNN–Transformer framework recovers a factorized, hierarchical representation of the fly’s chemosensory world in which modality, internal state, and valence are encoded along near-orthogonal axes rather than as an entangled high-dimensional code.

### Latent-space capacity constrains task-relevant dimensionality of whole-brain representations

The hierarchical structure of the latent space raises a natural question: how much embedding capacity does the model require at each level of task complexity, and what does this architectural requirement reveal about the underlying dimensionality of the neural representations themselves?

We swept *d*_model_, the dimension of the representation passed to the linear classification head, across five values (4, 8, 16, 32, 64) and evaluated test accuracy across 50 independent training runs per task, reporting the median and interquartile range (Fig. 2d). We distinguish throughout between *architectural capacity* (the number of dimensions available to the model, a hyperparameter under model architecture design) and *intrinsic dimensionality* [6, 56] (the number of linearly separable axes needed to capture the task-relevant variance in the neural population activity). Because the linear classification head can only exploit structure that is linearly separable within the bottleneck, the dimensionality at which accuracy saturates provides an upper bound on the task-relevant linear dimensionality, conditional on the representational power of the upstream encoder. Conversely, a statistically significant accuracy gain when moving from *d*_model_ = *n* to *d*_model_ = 2*n* implies that some task-relevant structure is geometrically inaccessible at the lower capacity, establishing a lower bound at *d*_model_ = 2*n*.

For the 2-class and 6-class tasks, accuracy was already near ceiling at *d*_model_ = 4 and remained flat across the entire range, plateauing at approximately 92–93% and 93–94% respectively with no further gain at higher capacities. A four-dimensional bottleneck therefore suffices for the whole-brain signatures of metabolic state and sensory modality, consistent with the decomposition in Fig. S3a, where each factor contributes a single dominant displacement that is largely unaffected by the other, so that two directions account for most inter-class variance.

The 16-class task revealed a qualitatively different profile. Accuracy rose steeply from 63% at *d*_model_ = 4 to 77% at *d*_model_ = 8, then continued rising to approximately 79–80% at *d*_model_ = 16, beyond which further increases in capacity yielded no meaningful gain. This monotonic rise followed by saturation contrasts sharply with the flat profiles of the simpler tasks, indicating that simultaneously separating all combinations of state, modality, and valence demands substantially more embedding capacity than any single factor in isolation: The near-origin placement of the conflicting-valence conditions along the valence axis (Fig. S3b) indicates the finer margins the model must resolve at this level. The plateau above *d*_model_ = 16 indicates that, for this architecture and classification objective, increasing latent capacity beyond 16 dimensions yields no additional gain in task-relevant decoding performance.

### Whole-brain population dynamics decode metabolic state, modality, and valence with high fidelity

The geometric separability of the latent space (Fig. 2) and the saturation of accuracy beyond *d*_model_ = 16 together indicate that the encoder maps whole-brain dynamics onto a low-dimensional representation in which metabolic state, modality, and valence are linearly separable. To characterize the structure of this decoding in more detail, we examined per-class performance and the pattern of residual errors across all three tasks (Fig. 3; Table T2).

**Figure 3:**
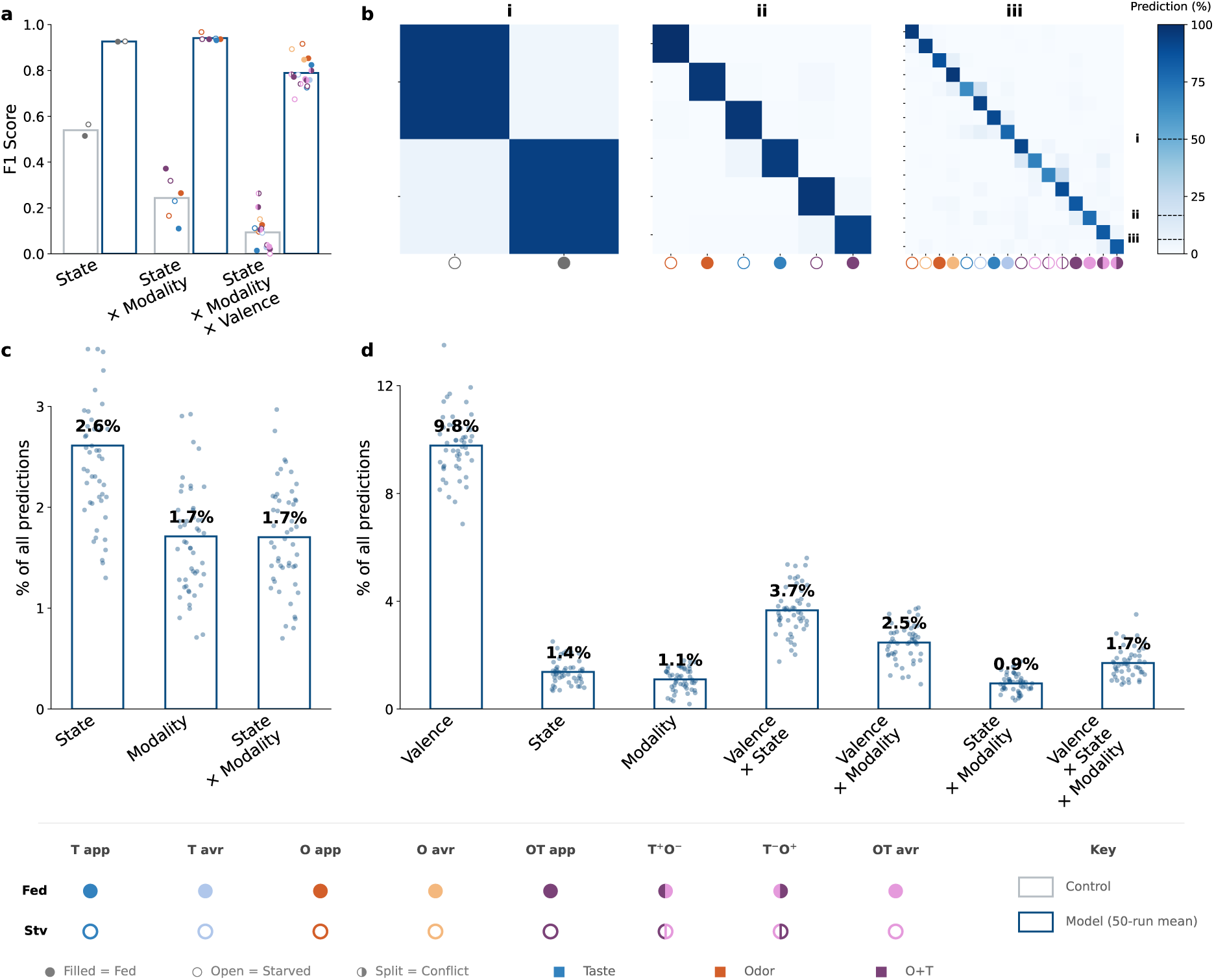
Whole-brain population activity supports high-fidelity decoding with structured residual errors. **(a)** Macro-averaged F1-score for the state (2-class), state–modality (6-class), and state–modality–valence (16-class) tasks. Bars indicate the mean across 50 trained models (blue) versus a single shuffled-label control (grey); dots show per-class F1. **(b)** Confusion matrices for the best model (*d*_model_ = 16) on tasks (i–iii); classes labeled as in the legend. Dotted lines on the shared colorbar indicate chance level for each task. **(c,d)** Errors decomposed by which experimental factor(s) differed between the true and predicted class: for the 6-class task (**c**: state, modality, or state × modality) and the 16-class task (**d**: state, modality, valence, and their pairwise and three-way combinations). Bars indicate the percentage of all test predictions in each category (mean across 50 trained models); dots show individual models.

The best models achieved 93.3%, 96.1%, and 84.2% test accuracy on the 2-class, 6-class, and 16-class tasks respectively, each substantially exceeding the corresponding shuffled-label control (54.1%, 27.8%, and 10%). Training was robust, with tight interquartile ranges across 50 independent runs (Table T2; Fig. S2). Notably, the 6-class task surpassed the 2-class task in absolute accuracy. A plausible explanation is that the 2-class model must find a single fed–starved boundary that holds across odor, taste, and combined conditions, whose whole-brain activity differs substantially; modality variation is then a nuisance the network must average over. Supplying modality labels separates these conditions in the latent space, so the fed–starved distinction need only be drawn within each modality, where it is more consistent.

Inspection of the confusion matrices and their factor-level decomposition revealed that residual errors were systematically structured rather than randomly distributed (Fig. 3b–d). In the 6-class task, state was the largest single error source: it was the only factor misclassified in 2.6% of predictions, against 1.7% for modality alone and 1.7% for both together (Fig. 3c). Counting both categories in which state was wrong, the 6-class model nonetheless identified metabolic state correctly in 95.7% of predictions, exceeding the 92.6% of the dedicated 2-class model.

In the 16-class task, this hierarchy reorganized: valence became the dominant source of confusion, accounting for 9.8% of all predictions in isolation, and a further 3.7% and 2.5% in combination with state and modality respectively (Fig. 3d). State’s error contribution dropped to a comparatively minor 1.4%. This error hierarchy parallels the latent geometry, in which valence is the axis with the finest displacement (Fig. 2iii, c; Fig. S3b).

### Spatial attribution and neuropil ablation analyses ground model decisions in distributed, anatomically plausible circuits

The factorized latent geometry established above shows what the model encodes; we next asked where in the brain this information originates. We approached this question at two complementary spatial scales: pixel-level attribution maps that highlight which parts of each image frame drive classification, and analyses of anatomically defined brain regions that quantify regional attribution and decoder reliance.

Grad-CAM++ attribution maps [57], computed at the first convolutional layer and averaged across all test samples per condition, revealed spatially structured attribution patterns that shifted systematically with experimental condition (Fig. 4a–b). Modality contrasts produced the most interpretable maps: odor conditions showed highest attribution in superior brain regions, taste conditions in more ventral regions, and combined stimulation produced a spatial union of both patterns. By contrast, attribution maps for state and valence contrasts were broadly distributed across much of the brain, making it difficult to isolate the spatial basis of these distinctions from attribution maps alone. This motivated a complementary, anatomically constrained analysis.

**Figure 4:**
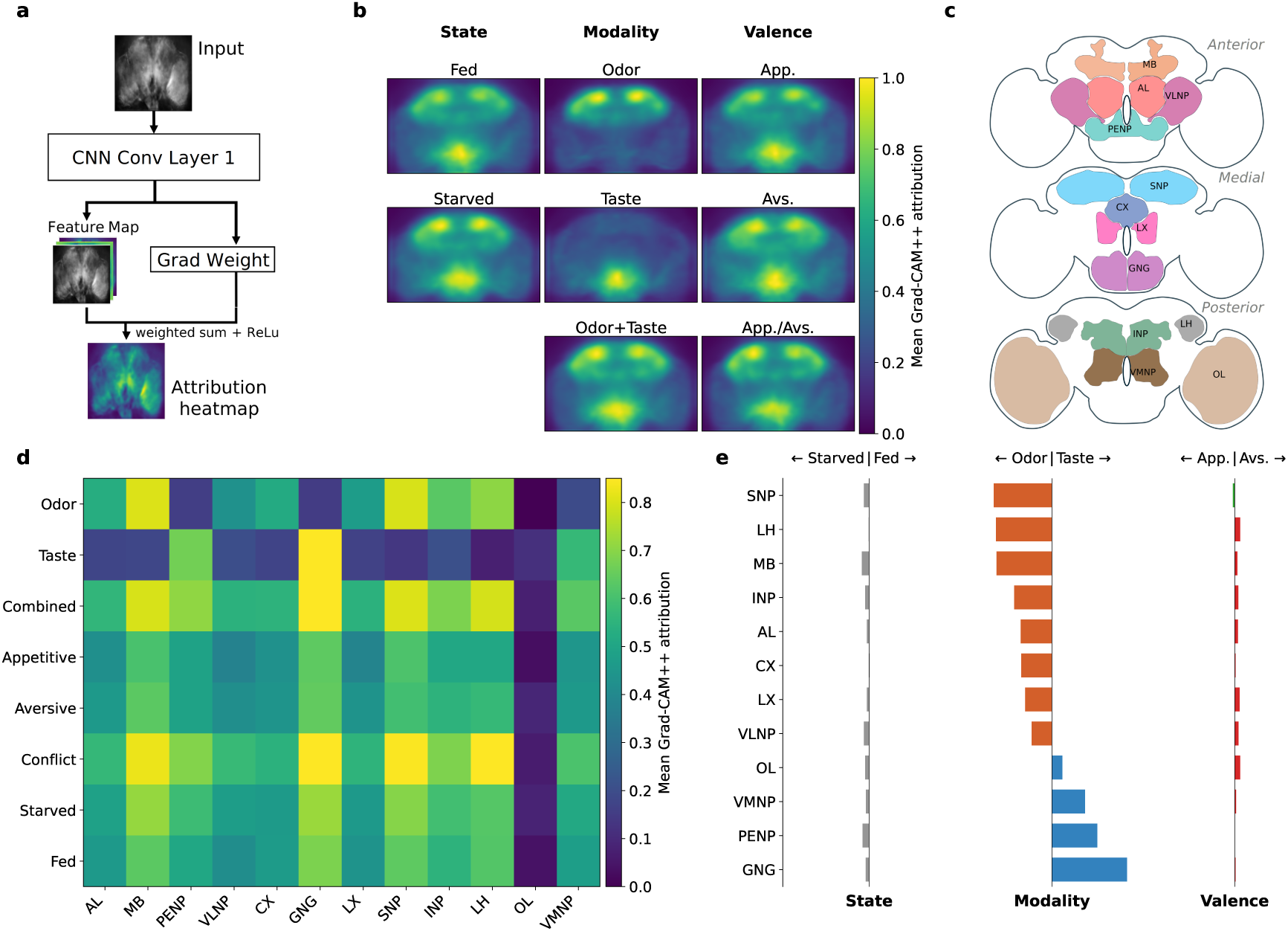
Spatial attribution reveals modality-specific neuropil patterns and distributed state and valence signals. **(a)** Schematic of the Grad-CAM++ spatial-attribution analysis applied to the first convolutional layer. **(b)** Grad-CAM++ attribution maps averaged across test samples for metabolic state (fed, starved), sensory modality (odor, taste, odor+taste), and stimulus valence (appetitive, aversive, conflicting). **(c)** Anatomical reference showing the 12 non-overlapping neuropil regions used for regional analysis in anterior, medial, and posterior views. **(d)** Mean Grad-CAM++ attribution within each neuropil mask, averaged across samples within the eight condition groups (odor, taste, odor+taste, appetitive, aversive, conflicting valence, starved, fed). **(e)** Pairwise differences in mean neuropil-level Grad-CAM++ attribution between opposing condition groups. Left: state (starved minus fed); middle: modality (odor minus taste); right: valence (appetitive minus aversive). Positive and negative values indicate greater attribution for the first or second condition of each contrast, respectively. Neuropil masks have zero voxel overlap in 3D.

To move beyond the qualitative attribution maps, we registered volumes to the JFRC2010 *Drosophila melanogaster* brain template and defined 12 anatomically distinct neuropil masks (Fig. 4c). For each test sequence, we computed the mean Grad-CAM++ attribution within each neuropil mask, yielding a group × neuropil attribution profile (Fig. 4d, see *Materials and Methods*). The mushroom body, lateral horn, and superior neuropils, three higher-order olfactory processing centers, were consistently among regions with highest attribution for odor conditions. For taste, attribution instead concentrated in the gnathal ganglia, the primary taste-processing center, and its neighboring periesophageal neuropils, to a lesser extent. The optic lobe yielded very low values throughout (dim blue colors), as expected for this peripheral visual region with no chemosensory role.

To test whether individual neuropils showed preferential attribution to one pole of each experimental factor, we computed pairwise differences in mean Grad-CAM++ attribution between the corresponding condition groups (Fig. 4e). For modality, odor–taste contrasts produced large positive or negative differences across multiple neuropils: olfactory-associated regions showed greater attribution for odor conditions, whereas the gnathal ganglia and neighbouring periesophageal neuropils showed greater attribution for taste. In contrast, the absolute attribution profiles showed substantial attribution across multiple neuropils for both fed and starved conditions and for both appetitive and aversive conditions (Fig. 4d). Accordingly, the corresponding fed–starved and appetitive–aversive differences remained close to zero across most neuropils (Fig. 4e). Thus, at the level of the neuropil categories analysed here, state and valence were broadly represented but showed little spatial segregation between opposing conditions. Virtual neuropil ablation provided convergent evidence (Fig. S4): several neuropils contributed to decoding state and valence in the absolute ablation profiles (Fig. S4a), whereas pairwise differences in ablation effects strongly distinguished odor from taste but were small for fed versus starved and appetitive versus aversive conditions (Fig. S4b). Together, these analyses indicate modality-specific specialization across neuropils, whereas state- and valence-related information is distributed across neuropils without strong segregation between opposing conditions.

## Discussion

Here, we show that deep representation learning applied to whole-brain volumetric calcium-imaging recordings reveals a factorized organization of neural population activity across metabolic state, sensory modality, and valence. High decoding accuracy demonstrates that mesoscale whole-brain activity contains sufficient information to distinguish these biological variables, but the central finding is that the learned latent space organizes whole-brain population activity into condition-specific clusters whose geometry is factorized along approximately independent dimensions (Fig. 2b,c; Fig. S3), despite the absence of explicit disentanglement constraints. The reproducibility of this organization across independent training runs further suggests that the observed factorization reflects structure in the underlying neural population activity rather than a specific solution of the optimization procedure. Importantly, these latent representations are biologically interpretable: spatial attribution and brain-region ablation analyses reveal anatomically structured patterns associated with each factor, linking latent dimensions to underlying neural populations and neuropil structures (Fig. 4; Fig. S4).

Neural populations across diverse species and brain regions often occupy low-dimensional manifolds in which task- and state-relevant variables are embedded within structured latent spaces [1, 2, 7]. Previous studies have revealed such low-dimensional organization in selected brain regions and behavioural contexts; our whole-brain representation-learning approach extends this perspective by uncovering a distributed population code spanning multiple sensory and internal variables simultaneously [24, 26, 58]. Task-informed approaches such as demixed PCA have further shown that mixed population activity can be decomposed into interpretable components associated with experimentally defined variables [new ref]. Here, however, the learned latent space organized brain-wide population activity into a factorized geometry even though the network received only flat class labels, with no access to the factorial structure or explicit disentanglement constraints (Fig. 2c; Fig. S3). The dimensionality required to decode these variables increased with representational complexity: compact embeddings were sufficient for simpler classification tasks, whereas representing the complete set of 16 experimental conditions required substantially greater latent capacity, with performance saturating around a 16-dimensional embedding (Figure 2d). Together, these findings indicate that whole-brain population activity can combine multiple task- and state-relevant variables within a structured but expandable representational space.

The structured geometry of the latent space also raises the question of whether whole-brain population representations may reflect computational principles related to attractor-based organization. Attractor-based models propose that neural populations can represent information through stable or metastable states in low-dimensional state spaces, providing a mechanism by which sensory inputs, internal variables, and memories can be transformed into robust representations [12, 15, 59]. In insects, attractor-like computations have been demonstrated mechanistically in specialized circuits, most prominently in the *Drosophila* central complex, where continuous ring-attractor dynamics support internal representations of orientation and navigation-related variables [16, 17, 60]. Complementary studies of insect olfactory circuits have revealed structured, low-dimensional population trajectories during sensory processing, illustrating that insect brains can represent information through distributed population dynamics beyond individual identified neurons [61, 62], with metastable and heteroclinic mechanisms proposed as possible dynamical explanations. The factorized geometry observed here shares conceptual similarities with such structured state spaces: whole-brain population activity is organized into separable dimensions capturing internal state, sensory modality, and valence. However, our analyses address the geometry of neural representations rather than their temporal evolution or causal stability; establishing attractor dynamics would require future measurements of state transitions, perturbations, and attractor basin structure.

The near-orthogonal factor axes recovered from whole-brain population activity reflect distinct biological dimensions that are known to shape insect behaviour. The modality axis corresponds to the anatomical segregation of olfactory and gustatory pathways, which enter the brain through distinct sensory channels before converging in higher-order regions [63]. In contrast, metabolic state and valence are known to be supported by distributed circuit mechanisms rather than a single anatomical locus. Hunger state is conveyed by broadly projecting neuromodulatory systems that reshape sensory processing and behavioural responses across multiple brain regions [38–40]. Similarly, valence information is represented across dopaminergic and downstream circuits involved in associative learning, value assignment, and behavioural selection [42, 45, 64, 65]. Consistent with this organization, spatial attribution and virtual neuropil ablation revealed a marked difference in regional organization across factors (Fig. 4; Fig. S4). Whereas olfactory and gustatory representations showed clear segregation across sensory-associated neuropils, fed versus starved and appetitive versus aversive conditions showed little segregation across the neuropil categories analysed here, despite distributed attribution and decoder reliance across multiple regions. This suggests that, at this anatomical scale, the opposing poles of state and valence are represented across many of the same brain regions rather than segregated into distinct sets of neuropils. The negligible attribution of the optic lobes further supports the specificity of the identified representations for internal and chemosensory variables rather than generic whole-brain activity patterns.

Our findings also highlight the potential of representation learning as a general strategy for analysing increasingly large-scale neural recordings [66]. By learning directly from whole-brain volumetric calcium-imaging data without prior cell segmentation, anatomical annotation, or region-of-interest selection, our framework preserves spatial organization while uncovering interpretable latent structure [67]. Such approaches may provide a scalable route for comparing brain-wide population dynamics across preparations and species, including whole-brain imaging approaches established in vertebrate models such as larval zebrafish and large-scale population imaging in rodents. Future extensions integrating functional imaging with concurrent behavioural measurements could enable joint representation learning across neural and behavioural spaces [51, 68], providing a route to test how latent population structure relates to action selection and adaptive behaviour.

While our approach reveals the geometry of whole-brain population representations, it does not yet resolve how these representations are dynamically generated or whether they play a causal role in adaptive behaviour. Future experiments combining brain-wide imaging with behavioural tracking and perturbations will be required to determine whether these latent representations influence action selection and behavioural variability. Similarly, integrating representation learning with cell-type-specific recordings, connectomic information [69], and targeted circuit manipulations will help identify the neuronal mechanisms through which distributed state- and valence-related representations emerge. More broadly, extending this framework across species and recording modalities may reveal whether factorized population representations constitute a general organizational principle by which biological brains organize diverse sensory and internal variables into flexible computational spaces.

## Materials and Methods

### Experimental Setup

#### Animals

We used 82 female *Drosophila melanogaster* (6–8 days post-eclosion) expressing the calcium indicator jGCaMP7s pan-neuronally (nSyb-GAL4 × UAS-jGCaMP7s; Bloomington Drosophila Stock Center, BDSC_51635 × BDSC_79032). Flies were reared on standard cornmeal medium at 25 °C under a 12 h : 12 h light–dark cycle. Animals were randomly assigned to fed (*n* = 41) or starved (*n* = 41) groups; starved flies were food-deprived for 18–24 h on a wet tissue prior to recording.

#### Calcium imaging

For imaging, each fly was head-fixed beneath a 25×/0.95 NA objective (Leica HC FLUOTAR L). Wings were removed to prevent flight movements; legs were left intact and free to move, with the exception of the front legs, which were immobilised to prevent interference with taste stimulus delivery. Whole-brain calcium activity was recorded using light-field microscopy (LFM) with widefield LED excitation (470 nm; Thorlabs M470L3) and a microlens array (MLA–S125-f12; RPC Photonics), following the acquisition and reconstruction pipeline of [58]. Volumetric Ca^2+^ signals were captured at 10 Hz with a native volume size of approximately 135 × 82 × 31 voxels, recorded by a scientific CMOS camera (Hamamatsu ORCA-Flash 4.0).

#### Stimulus protocol

Chemosensory stimuli were chosen to span appetitive, aversive, and conflicting valence categories within gustatory and olfactory modalities (Fig. 1c). Appetitive stimuli were 500 mM sucrose (taste) and 1% balsamic vinegar (odor); aversive stimuli were 4 mM quinine (taste) and 1% benzaldehyde (odor). Conflicting valence conditions paired an appetitive stimulus in one modality with an aversive stimulus in the other.

Taste stimuli (1 *µ*l) were delivered to the proboscis via a motorised glass capillary (1.5 mm outer diameter) for 5 s, followed by a 25 s inter-stimulus interval. Odorants were presented using a Syntech Stimulus Controller (CSS-55) through a tube positioned anterior to the head at a flow rate of 1000 ml min*^−^*^1^ for approximately 1 s, followed by an inter-stimulus interval of approximately 29 s. Each stimulus was delivered three times per recording session. Combined odor–taste stimuli were delivered simultaneously using both systems. The total session comprised three stimulus epochs separated by 30 s baseline periods (Fig. S1b).

### Data Preprocessing

#### Image processing and quality control

Raw light-field recordings were deconvolved to reconstruct volumetric brain activity [70] and motion-corrected using rigid-body volume registration (AFNI [71]). For each recording, a baseline frame was computed by averaging the first 25 s of each recording (a non-stimulus period) across time and z-planes. This baseline was subtracted from all subsequent frames to remove slow acquisition drift, with negative values clipped to zero.

We applied logarithmic transformation to the baseline-subtracted signal, computing log_10_(Δ*F* + *ε*) with *ε* = 10*^−^*^10^ for each voxel. We chose a log-transform rather than the conventional Δ*F/F*_0_ normalisation because the latter amplifies noise in regions with low baseline fluorescence, which is common in light-field reconstructions where out-of-focus light contributes non-uniformly across the volume. The log-transform stabilises variance and compresses the dynamic range while preserving the monotonic relationship between fluorescence and neural activity.

To reject frames dominated by motion artefacts or global fluorescence drops, we computed the mean log-intensity across the *x*–*y* plane for every *z*-slice at each time point and defined a global activity threshold as the recording-wide mean log-intensity. We retained frames in which *≥* 90% of z-planes exceeded this threshold. This procedure retained 443 *±* 68 frames per recording (mean *±* s.d.; approximately 37% of acquired frames). Supplementary (Fig. S1c) shows representative retained and excluded frames.

#### Volumetric data reduction and atlas registration

Given the limited number of recordings for full three-dimensional spatiotemporal learning, we compressed volumetric data along *z* dimension into two-dimensional summary images at each time step. We evaluated three projection strategies—mean, maximum, and variance across *z*—to identify the representation that preserved stimulus-evoked structure while suppressing depth-specific noise. In practice, mean-*z* projections yielded the most stable and visually interpretable images: they reduced the influence of single bright voxels (which dominated the maximum projection), while retaining more spatial structure than the variance projection, which overly emphasized fluorescence fluctuations. These mean-*z* frames then served as the base images from which we constructed the short image-sequences used for model input (sampling strategy detailed in Table T1). Based on these qualitative comparisons and the improved consistency of the resulting frames, we adopted mean-*z* projections as the basic imaging units for all subsequent analyses.

For the neuropil-based analyses (see below), corrected volumes were additionally registered to the standardized JFRC2018 template [72] via manual landmark registration [58]. Twelve non-overlapping neuropil supercategories, following the hierarchical brain nomenclature of [73], were used as regions of interest (Table T3, Fig. 4c). Each recording’s registration to the template was then used to bring the atlas neuropil boundaries into that recording’s own space, producing one mask per recording per neuropil, which were used in the subsequent Grad-CAM++ neuropil and virtual ablation analyses.

#### Data splitting

To prevent information leakage while making efficient use of the limited number of flies, we split the data at the level of entire stimulus epochs. Each recording session comprised three stimulus epochs separated by 30-s inter-epoch intervals. For each fly, two epochs were assigned to training and the remaining epoch was split evenly into validation (50%) and test (50%) subsets. This scheme ensured that no evaluation sequence shared time points with any training sequence, while allowing each fly to contribute to both training and evaluation without mixing temporal information across splits. The resulting split comprised approximately 50,000 training, 10,800 validation, and 10,800 test sequences across the full dataset, corresponding to approximately 4,500 sequences per condition. We tuned all hyperparameters exclusively on the validation set and report final metrics on the held-out test set, which remained untouched during model development.

### Model Architecture

We developed a hybrid CNN–Transformer architecture to learn representation and classify short LFM image sequences for three tasks of increasing complexity: (i) metabolic state (fed vs. starved; *K* = 2 classes), (ii) state × sensory modality (*K* = 6 classes), and (iii) state × modality × valence (*K* = 16 classes) (Fig. 1d). Each input sequence consisted of *S* = 5 consecutive mean-*z* frames of size 1 × 128 × 128 pixels (see Table T1).

#### Convolutional encoder

We processed each frame independently through three convolutional blocks, each comprising a 3 × 3 convolution (stride 2, padding 1) followed by ReLU activations. The first two blocks additionally included a 2 × 2 max-pooling layer (stride 2). Channel counts were 16, 32, and 64. An adaptive average-pooling layer reduced the output of final blocks to 64 × 1 × 1, which we projected to an embedding of dimension *E*_cnn_ = 16 via a linear layer.

#### Transformer encoder

We stacked the resulting per-frame embeddings into a sequence tensor of shape *S × E*_cnn_ and augmented them with learnable positional encodings. A single-layer Transformer encoder (2 attention heads; feedforward dimension 4 *× d*_model_; dropout *p* = 0.3) processed the sequence. We mean-pooled the encoder output across time to obtain a sequence-level representation of dimension *d*_model_.

#### Classification head

A single linear layer mapped the pooled representation to *K* class logits. The total model comprised approximately 28,000 trainable parameters (for *d*_model_ = 16; exact count varies slightly with *K*). A full list of architectural hyperparameters is provided in Table T1.

### Training Procedure

We trained all models using AdamW optimizer (learning rate 0.001, weight decay 10*^−^*^4^) for 1000 epochs with cross entropy loss and label smoothing (0.05 for 2-class; 0.1 for 6- and 16-class tasks). To quantify sensitivity to initialization, we repeated training with 50 random seeds per architecture–task combination. For each run, we retained the checkpoint with highest validation accuracy (no early stopping). We report mean *±* s.d. test performance across the 50 runs (Table T2).

### Evaluation

#### Baseline controls

To verify that classification performance reflected stimulus- or state-related information rather than recording-specific structure, we trained a label-permutation baseline. We used the identical architecture, input sequences, data splits, and training procedure, but randomly permuted class labels at the recording level: all sequences from a given recording received the same reassigned label, preserving class frequencies while destroying the correspondence between neural activity and experimental condition. We performed the permutation once with a fixed random seed and trained a single baseline model for each classification task. Validation and test labels remained unshuffled. Performance of this baseline therefore quantifies the extent to which the model can exploit recording-level confounds in the absence of meaningful label information.

#### Metrics

We assessed classification performance on the held-out test set using accuracy and macro-averaged F1-score. We used confusion matrices to characterise the structure of residual errors.

### Representation analysis

#### Latent space visualization

For each input sequence, we extracted the sequence-level embedding from the Transformer encoder output after temporal mean pooling and before the classification head. This vector summarizes spatiotemporal information across all frames. We projected embeddings from the held-out test set using t-SNE (perplexity = 30; scikit-learn [74]). t-SNE was used as a qualitative visualization tool to assess class separation and global structure in the learned embedding space; no quantitative analyses were based on t-SNE coordinates.

#### Factor-axis projections

To investigate the geometric relationships among experimental factors in the native high-dimensional latent space, we computed class centroids *µ_c_* = *|c|^−^*^1^ Σ*_i_*_∊*c*_*z_i_*for each of the *C* experimental conditions, where *z_i_*denotes the embedding of sample *i*. Centroid displacement vectors *v_c_*= *µ_c_ − µ^-^* were obtained by subtracting the global mean *µ^-^* = *N ^−^*^1^ Σ*_i_z_i_*.

Separately, we defined interpretable axes for each experimental factor by computing group-level means that pool all samples sharing a given factor level, regardless of other conditions. For instance, the state axis was defined as *a*^_state_ = (*z^-^*_fed_*−z^-^*_starved_)*/*|| *z^-^*_fed_ −*z^-^*_starved_, where *z^-^*_fed_ is the mean embedding across all fed samples irrespective of modality or valence. Modality and valence axes were constructed analogously from taste-versus-odor and appetitive-versus-aversive contrasts, respectively. Per-class centroid displacements were then projected onto these axes to visualize the arrangement of experimental conditions in the latent space (Fig. 2c).

Orthogonality between axes was assessed via cosine similarity, and the fraction of total centroid variance captured was computed as Σ*_k_*Σ*_c_*(*v_c_ · a*^*_k_*)^2^ */* Σ*_c_*||*v_c_*||^2^. Crucially, no orthogonalization was applied: the factor axes were used as computed from the data, and any observed orthogonality is therefore an emergent property of the learned representation.

### Spatial attribution maps

To identify image regions most influential for predictions, we generated Grad-CAM++ attribution maps [57] at the first convolutional layer (16 filters, 3×3 kernel, output 64×64). For each test sequence, we obtained Grad-CAM++ attribution per frame and averaged across the five time steps to yield a single attribution map per sequence. We then computed per-class maps by averaging across all sequences belonging to a given class. To facilitate visual comparison across conditions, we pooled per-class maps into eight factor-level maps by averaging across all classes sharing a given factor level (e.g. the “odor” map averages all four odor conditions regardless of state or valence). The resulting attribution maps were jointly min–max normalised to [0, 1] scale, so that relative intensity differences between conditions are preserved for visual comparison. To quantify neuropil-level attribution, we repeated this procedure on correctly classified test sequences only, upsampling each attribution map to 128×128 to match the resolution of the anatomical neuropil masks, allowing us to compute the mean Grad-CAM++ attribution within each masked region, yielding one attribution value per sequence per neuropil. These per-sequence values were averaged within the same eight condition groups used above (Odor, Taste, Combined, Appetitive, Aversive, Conflict, Starved, Fed) to obtain a group × neuropil attribution profile (Fig. 4d), and pairwise differences in attribution (starved−fed, odor−taste, appetitive−aversive) were computed from these group profiles (Fig. 4e).

#### Virtual neuropil ablation

To assess the anatomical specificity of the model’s learned representations, we used a virtual ablation approach. For each of the 12 neuropils, we generated a modified copy of the imaging dataset in which that neuropil’s voxels were replaced with a static baseline image, computed per recording as the mean neuropil signal over a pre-stimulus baseline window (frames 100–250) and held constant across all timepoints. We mean-projected the volumes using the same pipeline as the original data. For each of the 50 independently trained 16-class models, we computed per-class test accuracy on the unmodified data and on each of the 12 ablated datasets, yielding Δ accuracy = unmodified − ablated per class, neuropil, and model. Δ accuracy was averaged across the 50 models and normalised by each neuropil’s mean pixel footprint to control for differences in region size. Classes were pooled into the same eight condition groups used above, and pairwise differences in ablation effects were computed as starved *—* fed, odor − taste, and appetitive − aversive (Fig. S4).

### Implementation details

We implemented all models in PyTorch 2.4.1 (Python 3.12.3, CUDA 12.1) and trained them on a single NVIDIA RTX A6000 GPU (48 GB VRAM) running Ubuntu 24.04.2 LTS. Training a single model for 1000 epochs required 2.5 hours on average.

## Data Availability

Preprocessed whole-brain calcium-imaging datasets used for model training and evaluation, experimental metadata, trained model weights, and cached evaluation results sufficient to reproduce all figures reported in this study are available from GIN at https://gin.g-node.org/nawrotlab/DrosoEmbedding_WBCI and will be assigned a DOI upon publication.

The raw whole-brain calcium-imaging recordings will be deposited in a separate public repository and assigned a DOI upon publication.

## Code Availability

All custom code for preprocessing, model training, evaluation, and figure reproduction is available at https://github.com/NawrotLab/DrosoEmbedding. An archived, citable version of the code will be deposited in Zenodo and assigned a DOI upon publication.

## Supplementary Information

Whole-brain calcium activity was acquired by light-field microscopy from head-fixed flies during a 120 s protocol with three sequential stimulus presentations. Frames were retained for analysis only when global fluorescence remained stable across z-planes. Figure S1 illustrates the imaging setup, the stimulation timeline, and representative included and excluded frames across all eight condition groups.

**Figure S1:**
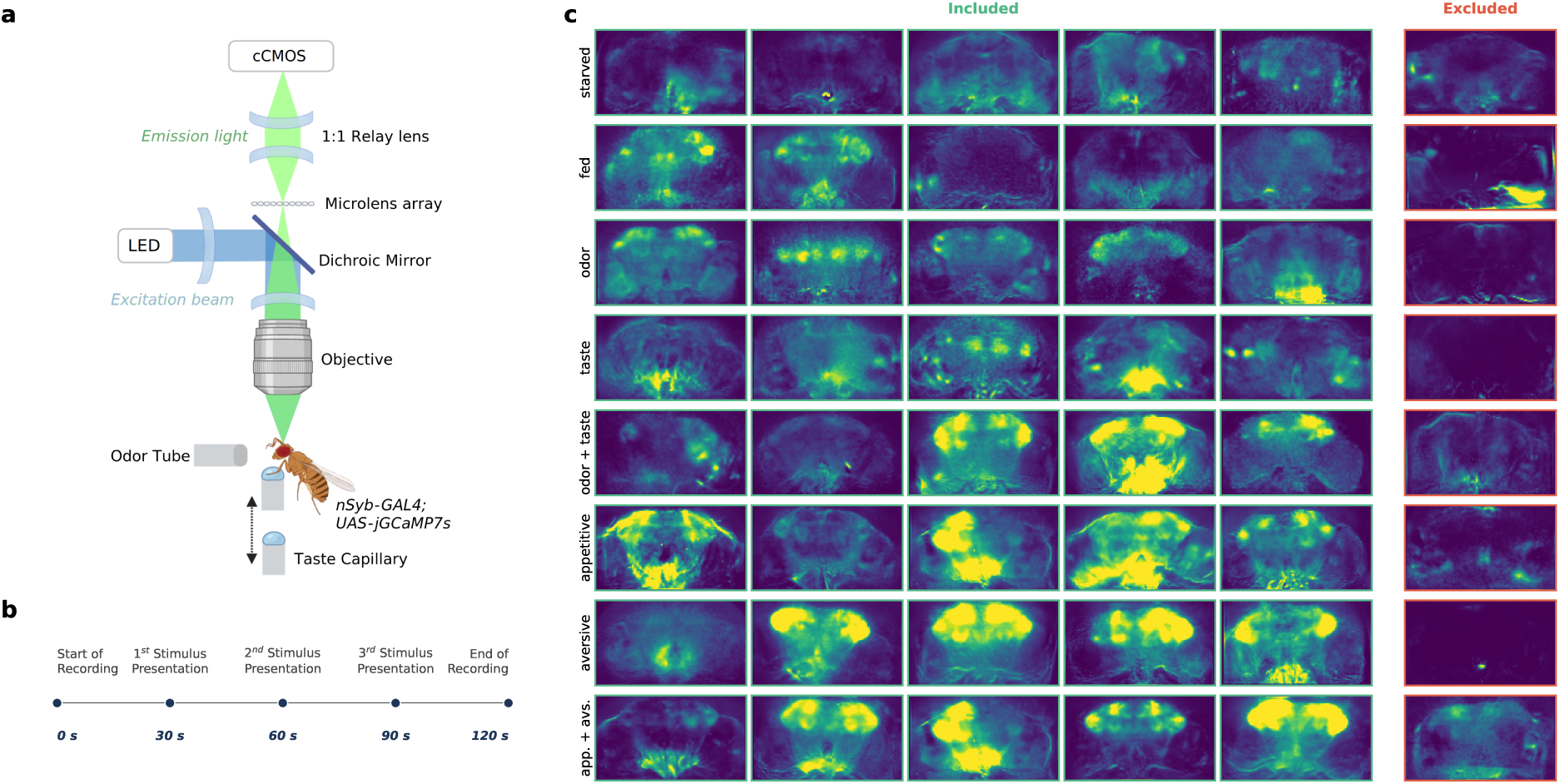
Imaging setup, stimulation protocol, and frame quality control. (**a**) LFM setup. Excitation light from an LED passes through a dichroic mirror and objective to illuminate the fly brain. Emitted fluorescence is collected through a microlens array and 1:1 relay lens onto a scientific CMOS (sCMOS) camera. The fly (*nSyb-GAL4; UAS-jGCaMP7s*) is positioned between an odor delivery tube and a taste capillary. (**b**) Recording timeline. Each recording spans 120 s, with three stimulus presentations at 30, 60, and 90 s. (**c**) Representative mean-intensity projections of whole-brain calcium activity, grouped by condition (rows): metabolic state (Starved, Fed), modality (Odor, Taste, Odor+Taste), and valence (Appetitive, Aversive, Conflicting). Each column within a row is drawn from a distinct recording to illustrate within-condition variability. Frames outlined in green passed quality control; frames outlined in red were rejected. Frames are individually normalised to the 1st–99th percentile of their source recording.

To confirm that training converged reliably and that reported accuracies reflect a stable optimum rather than a favorable snapshot, we tracked training and validation loss and accuracy across all 50 independent runs per task. Figure S2 shows the resulting curves for the 2-, 6-, and 16-class tasks.

**Figure S2:**
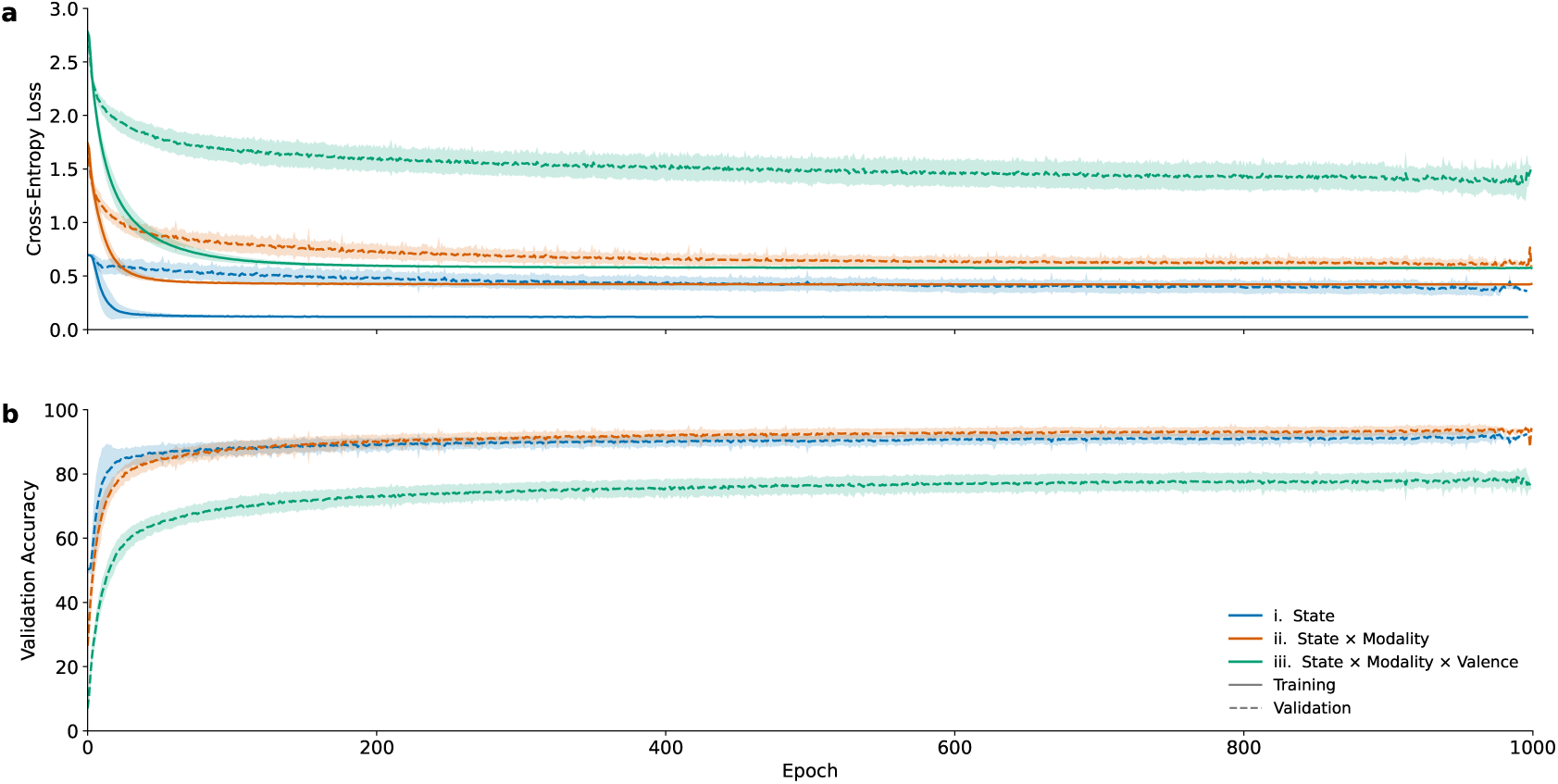
Training and validation curves across 50 independent model runs. (**a**) Cross-entropy loss and (**b**) validation accuracy over training epochs for three classification tasks: metabolic state (2-class, blue), state × modality (6-class, orange), and state × modality × valence (16-class, green). Solid and dashed lines show the mean training and validation curves, respectively; shaded bands indicate *±*1 standard deviation across 50 runs. All models were trained for up to 1000 epochs; the checkpoint with the highest validation accuracy was retained for evaluation (see table T1). Mean training length was 836 *±* 179, 834 *±* 231, and 884 *±* 138 epochs for the 2-, 6-, and 16-class tasks, respectively (min–max: 170–997, 165–1000, 337–1000); runs shorter than 500 epochs (*n ≤* 5 per task) are included up to their final recorded epoch. All three tasks converge with low inter-run variance across 50 random initialisations. Validation accuracy at convergence reached *∼*92%, *∼*93%, and *∼*78% for the 2-, 6-, and 16-class tasks, respectively.

The factor axes for state, modality, and valence were computed from group-mean embedding differences and applied without orthogonalisation (Materials and Methods). Projecting class centroids onto each axis in isolation, and onto each pair of axes, decomposes the combined projection of Fig. 2c into the contribution of each factor. Figure S3 shows these decompositions for the 6-class and 16-class models.

**Figure S3:**
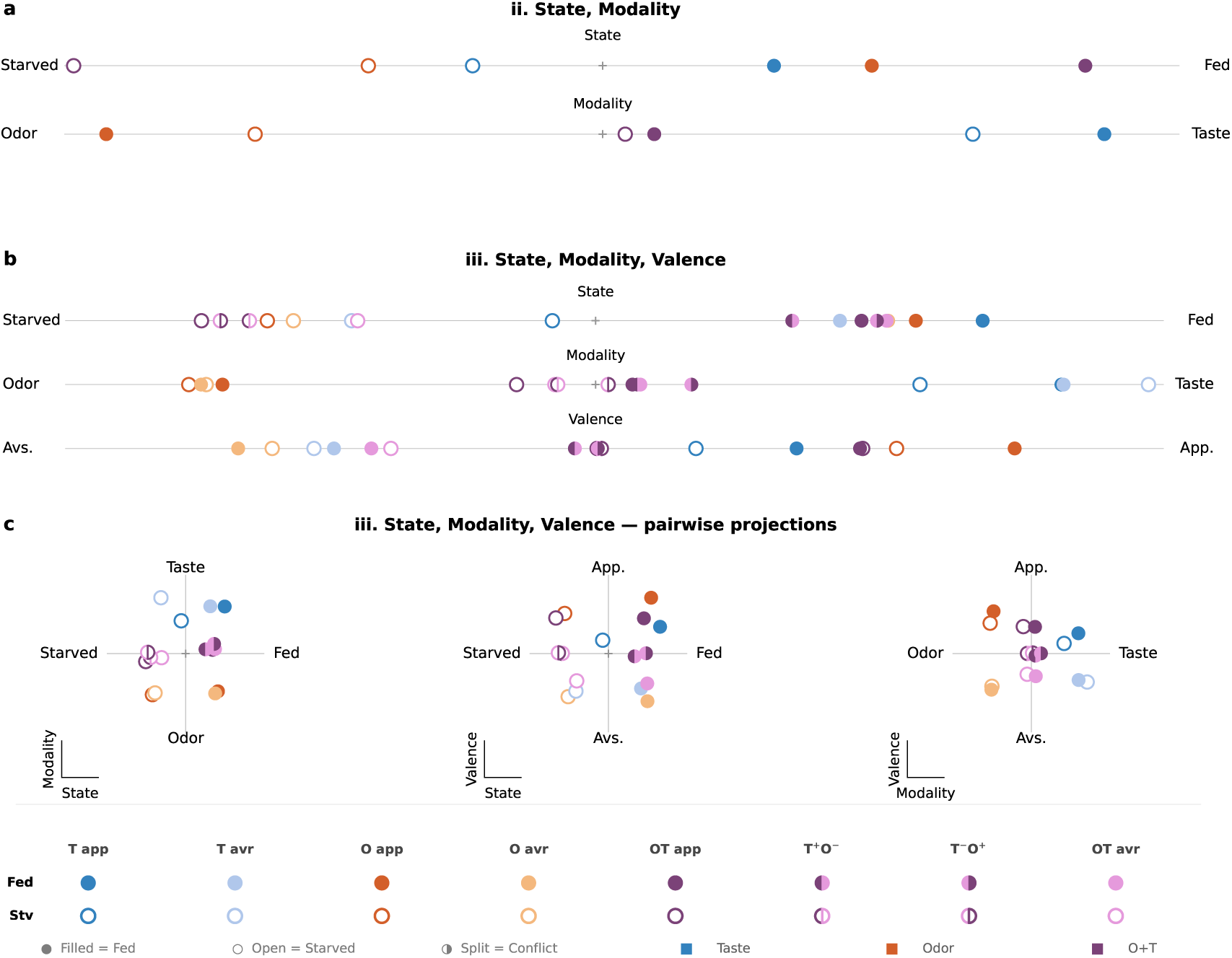
Latent-space separability of state, modality, and valence, individually and pairwise. Class centroids are projected onto the data-derived factor axes defined in *Materials and Methods*, decomposing the combined projections in Fig. 2c into their individual and pairwise components. **(a)** Task ii (State × Modality, 6-class model): centroids projected separately onto the State and Modality axes. **(b)** Task iii (State × Modality × Valence, 16-class model): centroids projected separately onto the State, Modality, and Valence axes. **(c)** Task iii centroids projected pairwise onto the State–Modality, State–Valence, and Modality–Valence axis combinations. Marker conventions as in Fig. 2: fill indicates state (filled = fed, open = starved, split = conflicting valence), color and shape indicate modality and valence.

To assess the anatomical specificity of the model’s learned representations, we used a virtual ablation approach. Each of the 12 annotated neuropils was virtually ablated independently by replacing its voxels with the per-recording mean ΔF/F over the pre-stimulus baseline window (frames 100–250), preserving resting-state spatial structure while removing stimulus-driven responses. The modified volumes were mean-projected and passed through the classifier. ΔAccuracy (unmodified minus ablated, in percentage points) was averaged across the 50 independently trained model checkpoints and normalised by neuropil size to account for differences in region footprint.

**Figure S4:**
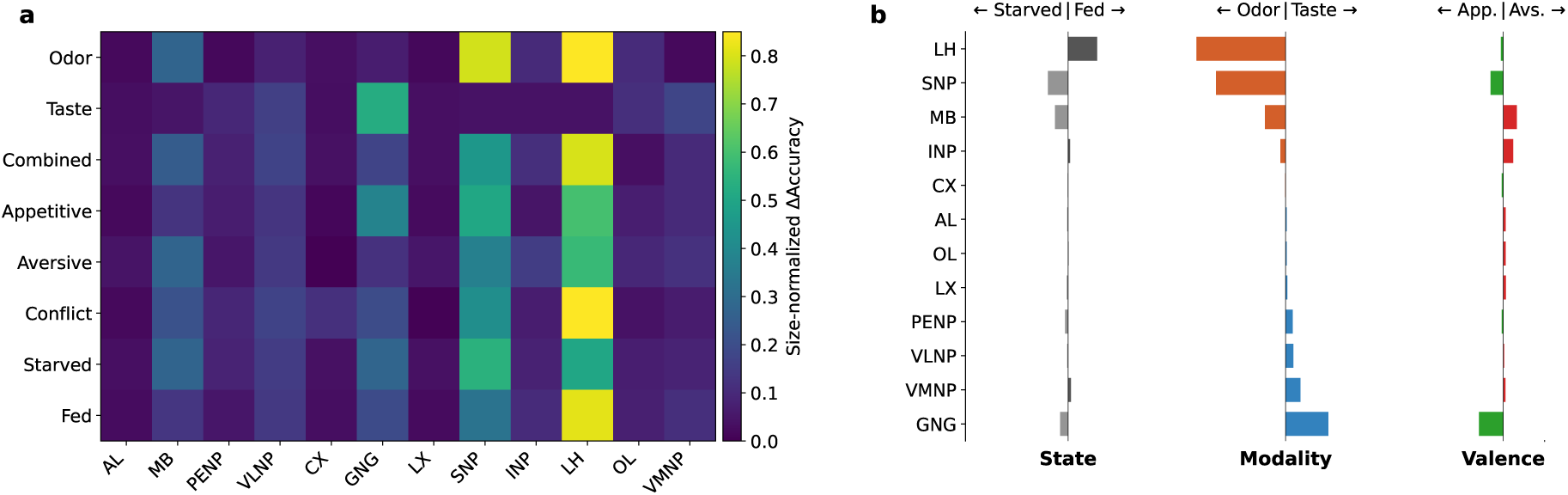
Region-specific decoding contributions quantified by virtual neuropil ablation. (**a**) Mean size-normalized change in classification accuracy (Δaccuracy; unmodified minus ablated) after virtual ablation of each neuropil, averaged across 50 independently trained models and grouped by experimental condition (rows: odor, taste, odor+taste, appetitive, aversive, conflicting valence, starved, fed; columns: 12 neuropils). Larger values indicate greater decoder reliance on the corresponding neuropil for that condition group. (**b**) Pairwise differences in virtual-ablation effects between opposing condition groups: state (starved minus fed), modality (odor minus taste), and valence (appetitive minus aversive). Bars show the difference in mean size-normalized Δaccuracy; positive and negative values indicate greater decoder reliance for the first or second condition of each contrast, respectively.

All models shared a single architecture and training procedure; only the transformer hidden dimension *d*_model_ varied (Table T1).

**Table T1:**
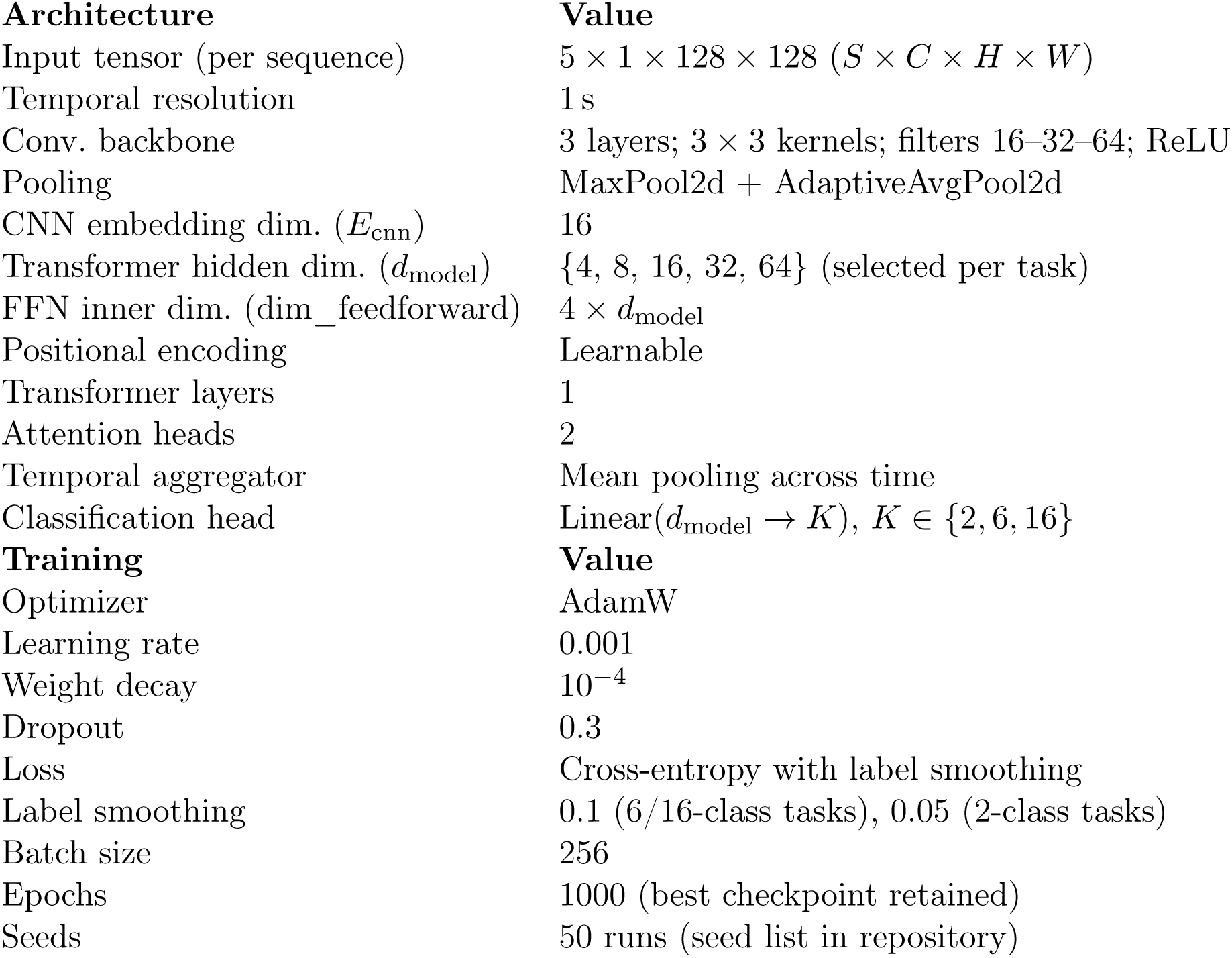
Model architecture and training hyperparameters. Hyperparameters held fixed across all task and dimensionality combinations. The transformer hidden dimension *d*_model_ was varied across *{*4, 8, 16, 32, 64*}*. The value selected per task by validation accuracy is marked with a yellow star in Fig. 2d. No learning-rate scheduling, gradient clipping, or mixed precision was used. The full seed list is provided in the code repository.

| **Architecture** | **Value** |
| --- | --- |
| Input tensor (per sequence) | $5 \times 1 \times 128 \times 128$ ($S \times C \times H \times W$) |
| Temporal resolution | 1 s |
| Conv. backbone | 3 layers; $3 \times 3$ kernels; filters 16–32–64; ReLU |
| Pooling | MaxPool2d + AdaptiveAvgPool2d |
| CNN embedding dim. ($E_{\text{cnn}}$) | 16 |
| Transformer hidden dim. ($d_{\text{model}}$) | $\{4, 8, 16, 32, 64\}$ (selected per task) |
| FFN inner dim. (dim_feedforward) | $4 \times d_{\text{model}}$ |
| Positional encoding | Learnable |
| Transformer layers | 1 |
| Attention heads | 2 |
| Temporal aggregator | Mean pooling across time |
| Classification head | Linear($d_{\text{model}} \rightarrow K$), $K \in \{2, 6, 16\}$ |
| **Training** | **Value** |
| Optimizer | AdamW |
| Learning rate | 0.001 |
| Weight decay | $10^{-4}$ |
| Dropout | 0.3 |
| Loss | Cross-entropy with label smoothing |
| Label smoothing | 0.1 (6/16-class tasks), 0.05 (2-class tasks) |
| Batch size | 256 |
| Epochs | 1000 (best checkpoint retained) |
| Seeds | 50 runs (seed list in repository) |

Per-task accuracy statistics underlying the main-text accuracy claims are listed in Table T2.

**Table T2:** Classification performance. Test accuracy across 50 independent training runs at *d*_model_ = 16, reported per task as best run, mean, median, and interquartile range. The control column shows the architecture-matched shuffled-label baseline (Materials and Methods).

| Task | Control | Best run | Mean | Median | IQR (25–75%) |
| --- | --- | --- | --- | --- | --- |
| i. State (2-class) | 54.1% | 93.3% | 92.6% | 92.9% | 92.1–93.5% |
| ii. State, Modality (6-class) | 27.8% | 96.1% | 94.0% | 94.2% | 93.4–95.1% |
| iii. State, Modality, Valence (16-class) | 10.0% | 84.2% | 79.0% | 79.0% | 77.5–80.3% |

Throughout the regional attribution and ablation analyses (Fig. 4), neuropils are referenced by the abbreviations defined in Table T3.

**Table T3:** Neuropil supercategories used in the regional attribution and ablation analysis. Abbreviations and full names of the twelve non-overlapping regions, following the hierarchical brain nomenclature [73].

| **Abbreviation** | **Name** | **Abbreviation** | **Name** |
| --- | --- | --- | --- |
| AL | Antennal Lobe | MB | Mushroom Body |
| CX | Central Complex | OL | Optic Lobe |
| GNG | Gnathal Ganglia | PENP | Periesophageal Neuropils |
| INP | Inferior Neuropils | SNP | Superior Neuropils |
| LH | Lateral Horn | VLNP | Ventrolateral Neuropils |
| LX | Lateral Complex | VMNP | Ventromedial Neuropils |

## Acknowledgments

We thank Martina Canova and Oğulcan Cingiler for their comments and feedback on the manuscript.

## Author Contribution Statement

Conceptualization: I.C.G.K., M.P.N.,V.R. Experimental design: P.B.,I.C.G.K. In vivo imaging: P.B. Data curation: A.A., K.Y.C. Model design for deep representation learning: A.A.,V.R. Design of spatial attribution and neuropil ablation analyses: A.A., V.R. Model training and Data analysis: A.A. Public release of code and data: A.A.,V.R. Supervision: V.R. Manuscript writing: A.A., M.P.N., V.R. Funding acquisition: I.C.G.K., M.P.N. All authors discussed the results and approved the final manuscript. Within each contribution, authors are listed in the order in which they appear in the author list.

